# Controlling *E. coli* gene expression noise

**DOI:** 10.1101/013797

**Authors:** Kyung Hyuk Kim, Kiri Choi, Bryan Bartley, Herbert M. Sauro

## Abstract

Intracellular protein copy numbers show significant cell-to-cell variability within an isogenic population due to the random nature of biological reactions. Here we show how the variability in copy number can be controlled by perturbing gene expression. Depending on the genetic network and host, different perturbations can be applied to control variability. To understand more fully how noise propagates and behaves in biochemical networks we developed stochastic control analysis (SCA) which is a sensitivity-based analysis framework for the study of noise control. Here we apply SCA to synthetic gene expression systems encoded on plasmids that are transformed into *Escherichia coli*. We show that (1) dual control of transcription and translation efficiencies provides the most efficient way of noise-vs.-mean control. (2) The expressed proteins follow the gamma distribution function as found in chromosomal proteins. (3) One of the major sources of noise, leading to the cell-to-cell variability in protein copy numbers, is related to bursty translation. (4) By taking into account stochastic fluctuations in autofluorescence, the correct scaling relationship between the noise and mean levels of the protein copy numbers was recovered for the case of weak fluorescence signals.

## I. INTRODUCTION

CELL-TO-CELL variability in protein copy numbers within isogenic populations are typically observed in various types of cells due to underlying random biochemical reaction processes [1], [2], [3]. The variability can lead to noise-induced cellular phenotypes such as cellular differentiation [3], multiple stability [4], and either sensitivity enhancement or suppression [5], [6]. Here we investigate the ability to differentially control the noise and mean levels of gene expression in *E. coli*.

Such differential control in gene expression has been achieved in different organisms such as yeast [7], [8], [9], soil bacteria [10], and mammalian cell lines [11]. In *E. coli*, systematic noise control have not been performed by perturbing promoter DNA sequences and ribosome binding sites, while most studies have been focused on genome-wide expression without such perturbations [16], [21], [2]. Here we aim to understand differential control of mean and noise levels of protein concentrations at the single cell levels of *E. coli*.

The approach we use is based on stochastic control analysis (SCA)[13], [12], a body of theory we developed and reported in previous publications. SCA is a sensitivity analysis framework, that is a direct extension of metabolic control analysis [14], [15] to the stochastic regime [13]. This approach is based on a *local* sensitivity analysis that can be applied to study first-order effects of finite-size perturbations. SCA can identify which parameters in stochastic systems – here, gene regulatory circuits – need to be varied by how much to achieve a desired control aim. This includes orthogonal control of noise levels with respect to mean levels, and simultaneous changes in noise and mean levels in the same or opposite directions for the same or different protein species. SCA can provide control efficiency and strength to identify the most effective control schemes that are experimentally relevant [12]. Here, we apply SCA experimentally to *E. coli* genetic systems.

In this paper, gene circuits are encoded on plasmid backbones, which are transformed into *E. coli* MG1655. The circuits express green fluorescent proteins (GFP) under the *lac*-promoter. We perturbed the expression system by inducing the promoter with isopropyl *β*-D-1-thiogalactopyranoside (IPTG) and using a library of ribosome binding sites (RBS). We found that by taking into account stochastic fluctuations in autofluorescence, scaling relationship between GFP signal noise and mean levels can be extended to weak signal regions, where autofluorescence becomes moderately strong. This implies that when fluorescent signals are not strong enough compared to autofluorescence, stochasticity in autofluorescence can be systematically taken into account to characterize cellular systems. In addition, we aimed to understand what the major sources of GFP signal noise are by investigating the scaling relationship between the GFP signal noise and mean levels via promoter induction and RBS perturbation. We found that one of the major noise sources is bursty translation.

## II. STOCHASTIC CONTROL ANALYSIS: REVIEW

SCA [12], [13] is a local sensitivity analysis based on control coefficients, which are defined approximately as percentage change in a response signal (*y*) divided by the percentage change in a system parameter (*p*):

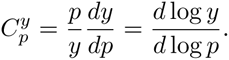

We note that the slope in the log-log plot of *y* vs. *p* corresponds to 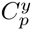. The response signal can be the mean or noise levels of mRNAs or proteins. The parameters can include transcription and translation efficiencies, degradation rates of mRNAs and proteins, dilution rate due to cell growth, and reaction rates of transcription-factor binding and unbinding from promoter regions, etc. Another important quantity in SCA, is the control vector, each element of which corresponds to a control coefficient for a given response signal (*y*):

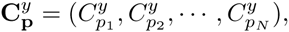

where *N* defines the number of parameters (dimension of the parameter space) that will be varied to control the value of *y*. In this paper, we are mostly interested in dual control of transcription and translation efficiencies, i.e., *N* = 2. One of the important properties of the control vector is that its innerproduct with a parameter perturbation vector *δ***p** becomes the amount of change in the response signal *dy*,

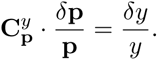

We can quantify which parameter value, and by how much it should be controlled to achieve specific control aims. For example, consider a case where the noise level of a protein needs to be reduced by 9%, while its mean level should remain the same. Here, the noise level (*n*) is defined by the variance divided by the squared mean value, i.e., squared coefficient of variation. Two control vectors for the noise and mean levels, 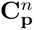 and 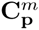, need to be computed based on a given mathematical model. System parameters need to be perturbed while satisfying

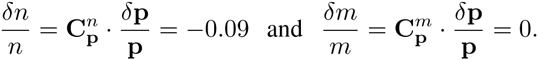

The perturbation vector *d***p***/***p** satisfying these two equations can be solved, but the solutions can be infinite. In that case, it is important to select the optimal control scheme (perturbation vector) among the possible solutions. For this, the control efficiency and strength were introduced [12]. Based on these two quantities, one can choose desired control schemes that are appropriate to systems of interest with the maximum control strength and/or efficiency.

## III. SCA FOR A SINGLE GENE EXPRESSION CASSETTE

We constructed plasmid expression systems that express green fluorescent protein (GFP) under *lac*-promoters in *E. coli* (Fig. 1A). The plasmid copy number in a single cell fluctuates in time because a set of plasmids are randomly partitioned during cell division and are synthesized in a stochastic fashion. Thus, the copy number of *lac*-promoters per cell fluctuates. For simplicity, we will assume that the plasmid copy number is tightly controlled, i.e., constant at the first level of approximation. The total number of *plac* will be the sum of the number of inactive and active *lac*-promoters (Fig. 1B), which will be set to a constant, *N_p_*. We call this the two-state model. The plasmid backbone that we used is pGA3K3 with the replication origin, p15A (*N_p_* =10 – 30). Based on this two-state model, we computed control vectors for the mean and noise levels of GFP fluorescence as shown in Fig. 1C (refer to the Materials and Methods and [12] for the control vector computation).

**Fig. 1.**
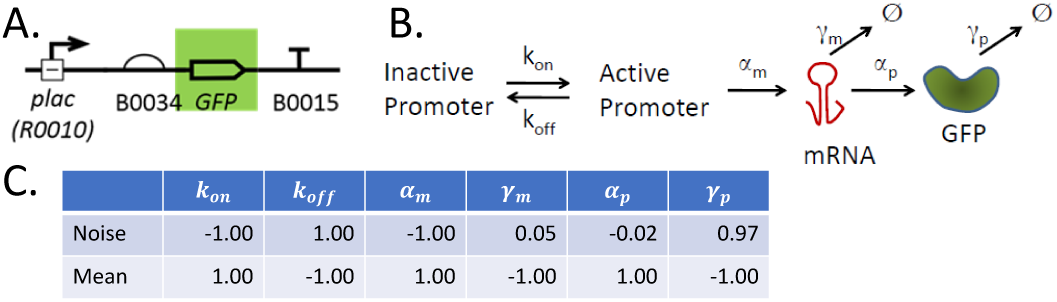
GFP expression system: (A) GFP is expressed under the *lac*-promoter (BioBrick part BBa R0010) with a ribosome binding site (BBa B0034). This expression cassette was placed in a low-medium copy number plasmid backbone (pGA3K3; origin of replication p15A). The ‘T’ symbol represents a terminator (double terminator used here to ensure transcription termination). (B) Two-promoter-state model. When the promoter is active, mRNA is transcribed with a rate constant *α_m_*. From the transcript, GFP is translated with a rate constant *α_p_*. mRNA and GFP degrade or are diluted with net rate constants *γ_m_* and *γ_p_*, respectively. (C) Control coefficients for noise and mean levels are listed. All control coefficients in the same row add up to zero (up to rounding error), satisfying summation theorems in SCA [13]. Parameters (unit: hr−1): *k_on_* = 50, *k_off_* = 51000, *α_m_* = 160, *γ_m_* = 30, *α_p_* = 1400, and *γ_p_* = 1 (refer to the Material and Methods for the detailed description of the mathematical model and its parameters).

The computed control coefficients show that noise can be controlled efficiently by varying *k_on_*, *k_off_*, *α_m_* or *γ_p_*; 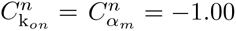, 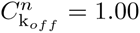 and 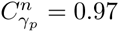, indicating that, for example, with an increase in *α_m_* by 10%, *n* will reduce by 10% (this is a first-order approximation, because control coefficients are defined locally). Similarly, with an increase in *γ_p_* by 10%, *n* will increase by 9.7%. For the mean level (*m*), any model parameter will efficiently change *m*, because the absolute values of all the control coefficients for *m* are equal to one.

## IV. MEAN LEVEL CONTROL

From the computed control coefficients, the mean protein levels can be controlled without changing the noise level (with a minor change, ~10 folds less than the change in the mean levels) by varying either *α_p_* or *γ_m_*. To confirm this theoretical prediction, we changed the translation efficiency *α_p_* by using a library of both ribosome binding sites (RBSs) and spacer sequences as shown in Fig. 2. Among them, four different spacers – TACTAG, AAAAAA=(A)6, (A)10, and (A)13 – that are placed between B0034 and the start codon showed distinct GFP expression levels when *plac* is fully active ([IPTG] = 1 mM). Here, the introduced spacer sequences are presumed to change ribosome binding affinity, in particular, translation initiation – the limiting step for a translation rate [23], [24]. We note that strong RBSs can recruit many ribosomes to mRNAs, causing an implication depending on the availability of ribosomes. This can apply an upper limit in the value of *α_p_*.

**Fig. 2.**
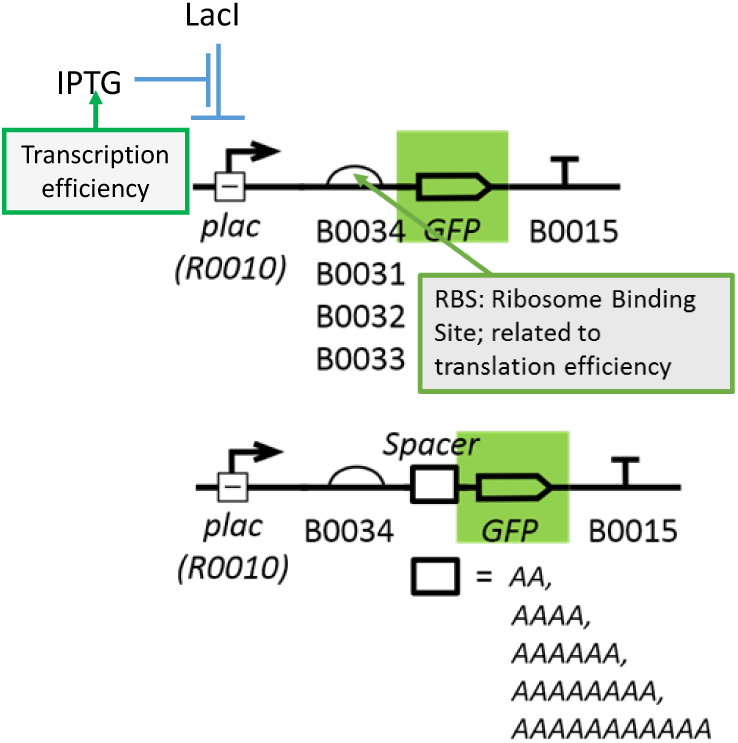
Perturbations in the GFP expression systems: The GFP expression cassette is placed in the plasmid backbone pGA3K3 in *E.coli* MG1655Z1 that constitutively expresses LacI. IPTG concentrations were varied for a given complex of ribosome binding site and spacer. ∼10 fold increase in the GFP noise level can be achieved without changing its mean level.

Based on our flow cytometry data, the mean level was successfully varied by using different spacer sequences as shown in Fig. 3 and Fig. 4A. We compared three different cases: Points A, B, and C in Fig. 3, corresponding to [IPTG]=1 mM. As shown in Fig. 4A, the rescaled probability density functions (pdfs) were overlapped with a minor discrepancy. This scale invariance confirms that the noise levels of all the points are the same.

**Fig. 3.**
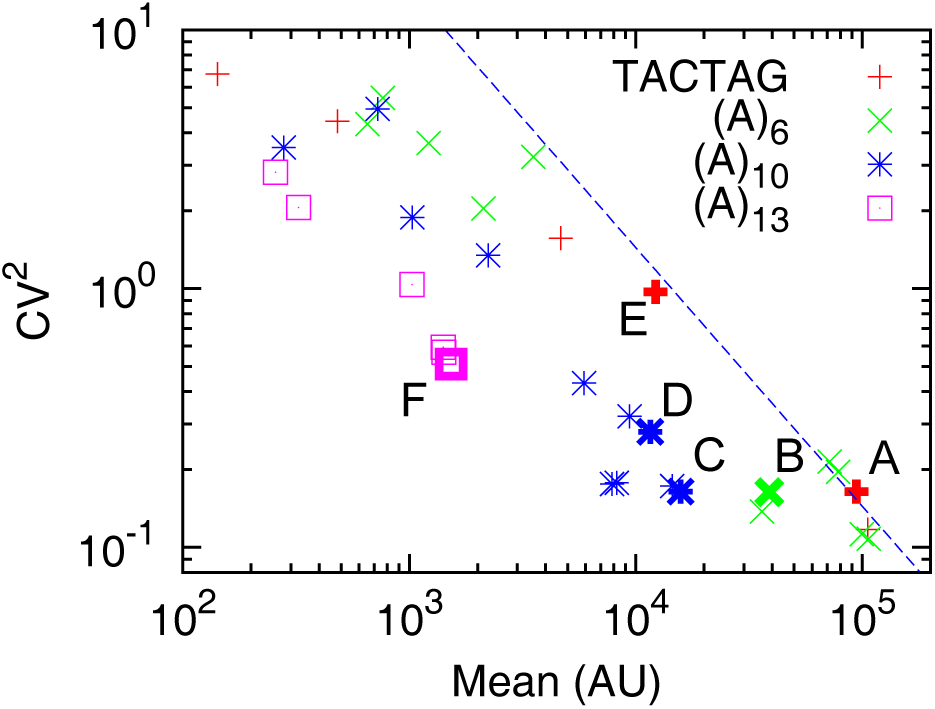
Scaling relationship between noise and mean levels: Four different cases of spacer sequences between the ribosome binding site Bba_B0034 and the start codon are shown. The same symbol represents the same spacer with different [IPTG]. The noise levels (squared coefficient of variation) are inversely proportional to the mean level. For comparison, a line with a slope –1 is drawn. The contribution to the noise level by background fluorescence signals was removed via the noise level correction method (Materials and Methods). For the IPTG concentration information, we refer to the Supplementary Notes Fig. 2.

**Fig. 4.**
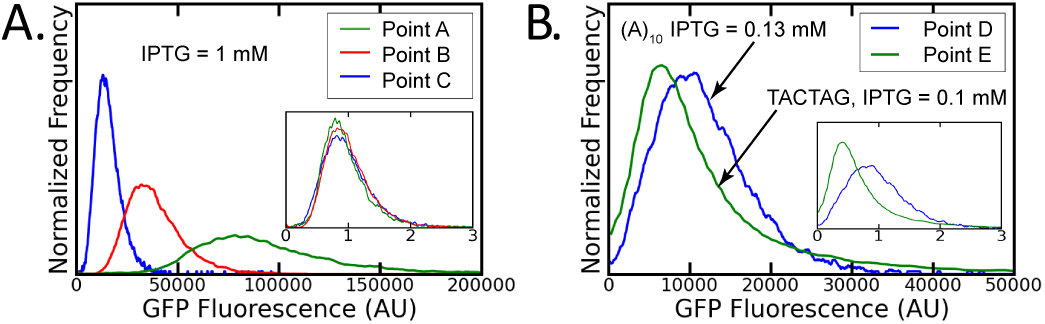
Probability density functions (pdfs) of GFP fluorescence signals measured from a flow cytometer: (A) Orthogonal mean level control: Points A, B, and C in Fig. 3 correspond to [IPTG]=1 mM. In the inset plot, both the pdfs were re-scaled by the mean values of their respective GFP fluorescence signals, so that the transformed pdfs are centered around one. (B) Orthogonal noise level control: Points D and E. Both the [IPTG] and the spacer sequences were varied. [IPTG]=.13 mM for Point D and .1 mM for Point E. Autofluorescence was removed via the fluorescence histogram correction method (Materials and Methods).

*Scale invariance in the gamma distribution*: Furthermore, the observed invariance implies a special property that we need to consider carefully. This invariance property is satisfied by the gamma distribution function as shown in the Materials and Method section when the burst size is rescaled together. This implies that the difference between the system parameters of Points A, B, and C is only the burst size. For these Points, different spacer sequences were used between B0034 and the start codon, while the *lac*-promoter was fully induced (saturated). Thus, the translation rate constant *α_p_* is expected to be varied for these three Points and the burst size must be closely related to *α_p_*, which is consistent with theoretical prediction based on our model (Eq. (3)). This result supports that the observed distribution functions are the gamma distributions (confer to [25], [26] about claims for other types of distribution functions). To confirm this, we fit the GFP pdfs to the gamma distribution functions as shown in Fig. 5. We confirmed that the pdfs follow the gamma distributions well.

**Fig. 5.**
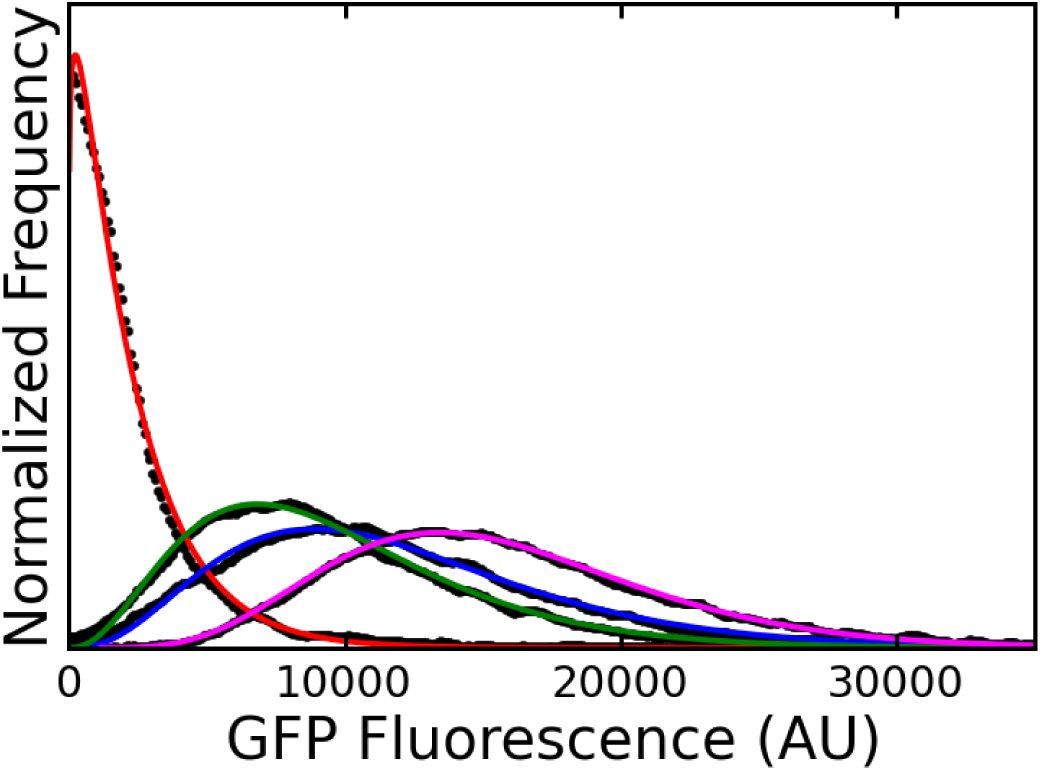
True GFP signal distribution function for the (A)_10_ cases with different IPTG concentrations: The fluorescence histogram correction was applied to remove autofluorescence effects. The true GFP signal distribution satisfies the Gamma distribution functions.

## V. NOISE LEVEL CONTROL

As discussed above, the noise level can be efficiently controlled by varying *k_on_*, *k_off_*, *α_m_* and *γ_p_*. However, when these parameters are changed, the mean level also changes with the same fold difference but in the opposite direction; for example, in Fig. 1C, 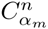 and 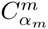 are −1.00 and 1.00, meaning that when *α_m_* is increased by x%, the noise level decreases by x%, while the mean level increases by x%. Thus, to change the noise level without changing the mean level, we must vary at least two different parameters simultaneously.

Since the mean level can be controlled almost independently of the noise level by changing *α_p_*, we will vary *α_p_* along with one of the parameters in {*k_on_, k_off_, α_m_*} to compensate for the change. The reason that we did not choose to vary *γ_p_* is that this parameter is highly dependent on cell growth rate, rather than protein degradation in *E. coli*; GFP lifetime is much longer than the cell doubling time ∼1 hr in M9 media.

Based on the SCA, an individual change in *k_on_*, *k_off_*, and *α_m_* and any combination of the individual changes can vary the noise and mean levels while satisfying the same scaling law: *n* = *c/m* with *c* a constant (not varied). We note that the ratio of control coefficients for *n* and *m* for a given parameter, e.g., *k_on_*, has a graphical meaning: In Fig. 3, when [IPTG] is varied, the corresponding data point shifts (e.g., Point C → D) and the slope of the shift in the loglog plot corresponds to the ratio of the control coefficients: 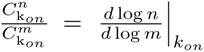. For *k*_*on*_, *k*_*off*_, and *α_m_* the ratios are the same: 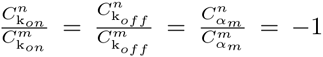. This implies that the directions of data point shifts in the log-log plot of *n* vs. *m* are identical for each individual perturbation of *k_on_*, *k_off_*, and *α_m_*, and thus for any combination of these three individual perturbations. Therefore, the shift of data points with the slope of 1, observed when varying IPTG concentrations as shown in Fig. 3, cannot determine which parameters among *k_on_*, *k_off_*, and *α_m_* were affected by IPTG concentration changes. We note that in [21] promoter perturbations in *E. coli* was claimed to affect *k_off_* only.

As shown in Fig. 3, by using a library of RBS as well as different concentrations of IPTG, the noise level was controlled and ∼10 fold change in the noise level was achieved without changing the mean level. What is the biological reason that noise can be increased in this way? In other words, what causes to increase the value of the Fano factor? In the scaling relationship, *n* = *c/m*, *c* is the Fano factor, which is expressed for the case of *E. coli* (Materials and Methods):

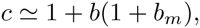

where

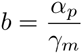

and

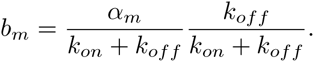

*b* is the translational burst size, quatifying the number of proteins that are synthesized from a single mRNA during the mRNA lifetime (1*/γ_m_*). *b_m_* is the transcriptional burst size, quantifying the number of mRNA that are synthesized per plasmid during the time-scale (1*/*(*k_on_* + *k_off_*)) of gene switching (refer to the Materials and Methods). The Fano factor depends on both transcriptional and translational bursts. In our case of lacI^*q*^ expression of LacI, we can neglect the transcriptional burst (Materials and Methods). Thus, the Fano factor becomes *c* = 1 + *b*, and *n* can be expressed as

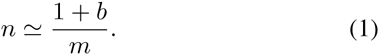

The Fano factor can be increased by applying stronger translation efficiencies (from Point C to A) and remains the same by decreasing [IPTG] (from Point A to E), leading to the increase in the noise level without changing the mean level.

The translational bursts lead to longer-tail pdfs, more precisely, higher cutoff values in the pdfs (in the gamma distribution, there is an exponential factor *e^-x/b^* and *b* acts as a cutoff value): Figure 4B shows that a longer tail in the GFP pdf can be generated by using stronger translation efficiency (Point D → Point E).

Another interpretation for the observed longer tail in Fig. 4B is that the major source of fluctuations in the protein copy numbers is in mRNA copy numbers, which merely get amplified by the translation rate in a non-bursty way:

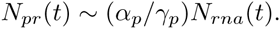

The variance in protein expression levels becomes

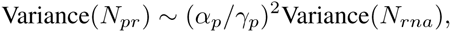

resulting in that the noise level does not depend on *α_p_*:

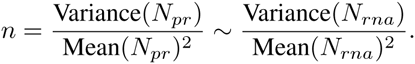

Since the protein mean level *m* must be proportional to the translation rate constant *α_p_* (i.e., *m* = *βα_p_* with *β* a constant), we obtain again similar scaling relationship:

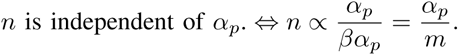

The Fano factor again increases with *α_p_*. Therefore, based on the scaling relationship alone, it is difficult to differentiate whether the translation is bursty or not.

## VI. TRANSLATION IS BURSTY

We claim that translation processes are bursty. The data points in bold in Fig. 6 correspond to different RBS strength but the same level of [IPTG] equal to 1 mM, where the *lac*-promoter becomes constitutively active. The noise values for TACTAG, (*A*)6, and (*A*)10 were similar, but the noise level for (*A*)13 was higher than the rest. This difference cannot be explained in the non-bursty translation scenario, because *n* should be independent of *α_p_*, i.e., RBS strength. In the bursty translation scenario, the noise level can be dependent on the value of *α_p_*, especially when the value of *α_p_* is similar to that of *γ_m_*. Since *m* is proportional to *α_p_* (*m* = *βα_p_*), Eq. (1) becomes

**Fig. 6.**
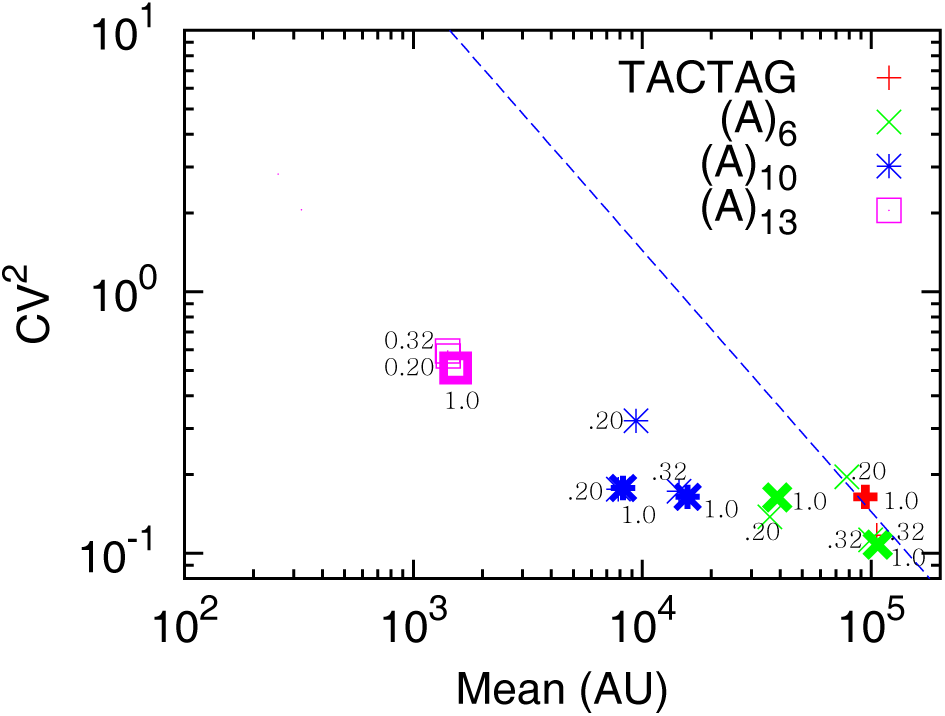
GFP mean and noise levels for the cases that *lac*-promoters are fully induced ([IPTG] = 0.20, 0.32, 1.0 mM). Two biological replicates were used for (*A*)6 and (*A*)10.

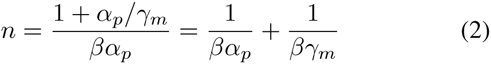

For a strong RBS such as the TACTAG case, *α_p_* can be roughly around 1400 hr^−1^ (Materials and Methods). In this case, *α_p_/γ_m_* ∼ 1400*/*30 ∼ 47, i.e. much larger than 1. Thus, Eq. (2) becomes 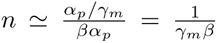, resuling in that *n* is independent of *α_p_*, which is what we observed from Point A to C. For the case of (*A*)13 (Point F), the RBS strength is reduced by ∼ 60 times (by comparing the mean levels of Point A and F) and *α_p_/γ_m_* ∼ 47*/*60 ∼ 0.8. Thus, *n* becomes dependent on *α_p_* (Eq. (2)). As *α_p_* decreases, *n* increases. This is consistent with our observation.

## VII. SCALING RELATIONSHIP BETWEEN THE NOISE AND MEAN LEVELS

Figure 7 shows that the scaling relationship *n* = *c/m* can be observed after autofluorescence was systematically removed, even for the small mean value region, where the autofluorescence interferes with the true GFP signals. We took into account the stochasticity in autofluorescence and assumed that the fluctuations in the autofluorescence signals are statistically independent of the true GFP signals. Under this assumption, we compensated for the autofluorescence effect in two different ways: (1) direct noise level correction (red squares in Fig. 7) and (2) fluorescence histogram correction (blue triangles in Fig. 7; an example is presented in Fig. 8) (Materials and Methods). This implies that it is important to take into account the stochasticity of the autofluorescence signals when characterizing the systems by using the pdfs of fluorescence signals.

**Fig. 7.**
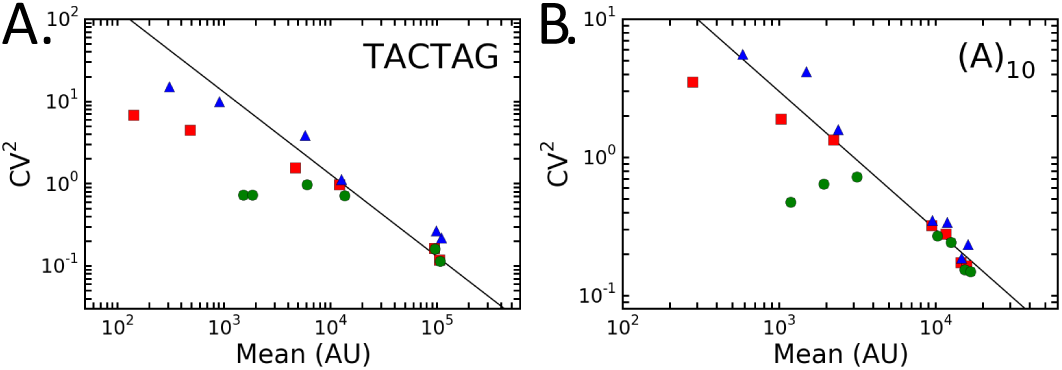
Effect of autofluorescence on scaling relationship between *n* and *m*: Autofluorescence was removed in two different ways via (1) the noise level correction method (red squares) and (2) the fluorescence histogram correction method (blue triangles) (refer to the Materials and Methods). To show the trend clearly, the data corresponding to the same biological replicate are only shown.

**Fig. 8.**
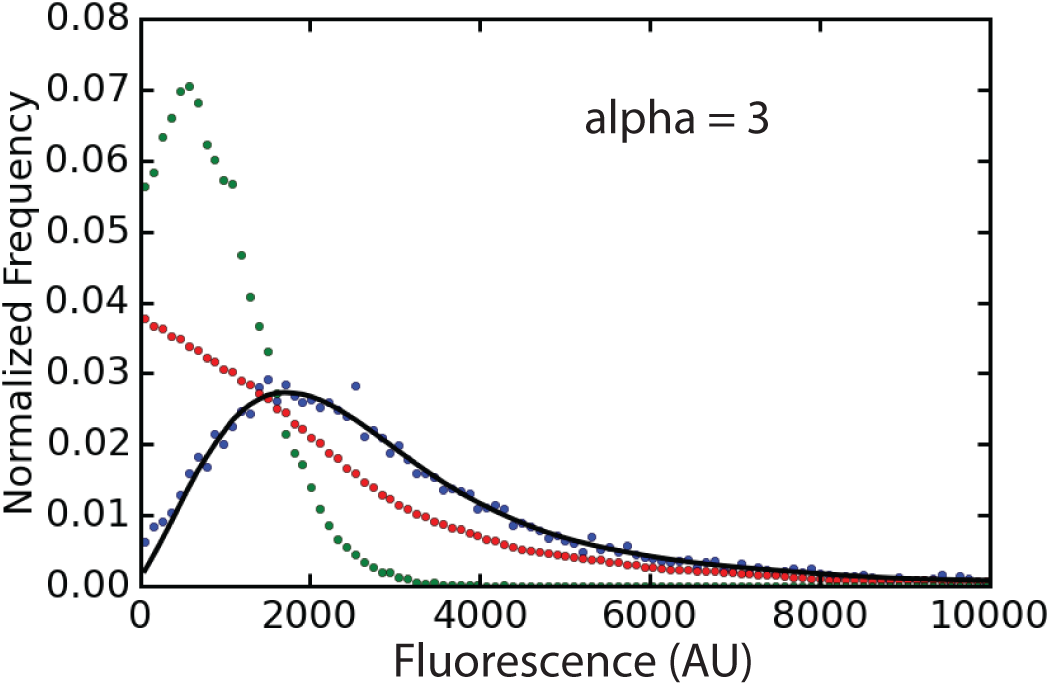
Autofluorescence compensation: The green dots correspond to autofluorescence (IPTG=0 case), the blue dots to the measured GFP signals, and the red dots to the optimized solution *S*, i.e., the pdf of the true GFP signals. The black line is to verify the optimized solution *S* can generate the measured GFP pdfs via convolution (refer to the Materials and Methods).

## VIII. CONCLUSIONS

In summary, we perturbed the strength of ribosome binding sites and investigated scaling relationship between the mean and noise levels of the expressed proteins. We confirmed that translational bursts are one of the important sources of noise at the protein level by using our numerical sensitivity analysis method, SCA, and the analytical structure of noise propagation. To investigate the scaling relationship further in detail, we compensated the effect of autofluorescence by taking into account stochasticity in the autofluorescence and recovered the expected scaling relationship even when autofluorescence becomes moderately strong. This shows that the autofluorescence can be systematically removed and its compensation can be applied to characterize cellular systems.

## MATERIALS AND METHODS

### A. GFP expression circuits and strains

All genetic components used in this manuscript are BioBrick parts, from which genetic circuits were constructed by using the Gibson assembly method [30]. The constructed circuits were integrated into a low-to-medium copy number plasmid pGA3K3 with a Kanamycin resistance gene and *Escherichia coli* MG1655 Z1 was transformed with the plasmids. The strain (lacI^*q*^) constitutively overexpresses LacI from its chromosome.

### B. Cell Growth and Flow Cytometry Measurements

*E. coli* strains were grown to OD600∼0.2 in 2 mL Luria-Bertani (LB) media (Becton Dickinson) with kanamycin 50 *µ*g/mL at 37°C and 300rpm in a shaker. The cultures were diluted 1:200 into 200 *µ*L prewarmed fresh M9 media (Teknova 2M1990) in 96-well plates (Costar 3904) with kanamycin 50 *µ*g/mL. 12 different IPTG concentrations (0 mM, 0.02~1 mM) were used for each well (refer to the Supplementary Notes for more detailed information on IPTG concentrations) and grown to OD600=0.3-0.4 in a shaker (37°C, 300 rpm). For the flow cytometry measurements, the grown cultures were diluted 1:4 in 1xPBS. A Sony Biotechnology ec800 flow cytometer was used with a 525 nm filter and a 488 nm excitation laser for GFP fluorescence. 100,000 events were collected for each sample and gated by using a 2-D normal distribution (Bioconductor flowCore norm2filter function with scale.factor=1) [31] within the R software environment as well as by using python package FlowCytometryTools (http://gorelab.bitbucket.org/flowcytometrytools/#). To prevent well-well contamination we executed a Medium Flush cycle after each sample well. When computing the mean and noise levels of GFP signals, background fluorescence was removed by using the mean and noise levels of GFP signals, or the signal histogram for the case without IPTG for each different gene circuit.

### C. Mathematical Model

A two-state model [27], [16], [28] is introduced to describe active and inactive states of a promoter along with transcription and translation processes:

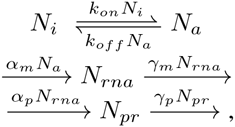

where *N_i_* denotes the number of inactive promoters, *N_a_* that of active promoters, *N_rna_* the RNA copy number, and *N_pr_* the protein copy number. All the above reaction events are generated stochastically. The noise level, *n*, of *N_pr_* can be analytically solved (refer to the Supplementary Notes of [27] for all the detailed derivation):

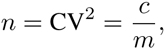

with *m* denoting the mean value of *N_pr_* and the Fano factor *c* is expressed as

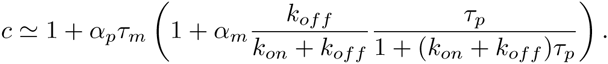

Here, we assumed that the protein lifetime *τ_p_*, defined by 1*/γ_p_*, is much larger than the mRNA lifetime *τ_m_*. We note that control coefficients for *n* can be computed from this equation analytically. We assume that the inactive and active promoter states switch back and forth many times during the protein lifetime (*k_on_* + *k_off_ ≫ γ_p_*). We believe this is the case of our experiments and refer to the parameter value estimation described below. In this case, the above equation can be further approximated:

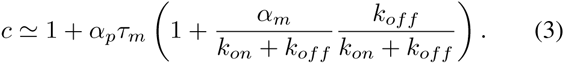

Here, 1*/*(*k_on_* + *k_off_*) is the time scale of the promoter state switching and *k_off_ /*(*k_on_* + *k_off_*) is a suppression weight because the promoter state follows the binomial distribution (due to the fact that the total promoter number is constant) instead of the Poisson distribution. Thus, the second term in the parenthesis can be considered as a transcriptional burst size *b_m_*. In our case, *α_m_/*(*k_on_* + *k_off_*) *∼* 160*/*(50 + 51000) ∼ 0.003. Thus, *c* can be further simplified:

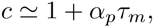

implying that the change in the IPTG concentration has no effect on the Fano factor, *c*, thus moving along the line of slope −1 as shown in Fig. 3.

#### D. Model parameter estimation

Transcription rate constant, *α_m_* = 160 hr^−1^: The *lac*-promoter strength, when fully induced with IPTG, was shown*∼*1.5 time stronger than J23101 by directly measuring the transcript levels with our malachite-green aptamer probes (refer to Figure 6.3 of [22]). For J23101, *α_m_* was estimated at 0.03 sec^−1^=110 hr^−1^ [32]. Thus, *α_m_* for our *lac*-promoter can be estimated at 160 hr^−1^.

We used the translation rate constant, *α_p_* = 1400 hr−1 (Supplementary Notes in [33]), the dilution rate, *γ_p_* = 1 hr^−1^, and the degradation rate constant of mRNA, *γ_m_* = 30 hr^−1^.

Gene inactivation, 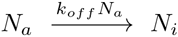 The number of the inactive promoters is denoted by *N_i_* and that of the active promoters, *N_a_*. The sum of *N_i_* and *N_a_* is equal to the copy number of the plasmids, *N_p_* (considering that one *lac*-promoter is included per plasmid). Here, we used *N_p_* 10 (http://parts.igem.org/Part:pSB3K3; pGA3K3 is a variant of pSB3K3). *k_off_* is related to the search time for LacI to find *lac*-promoter. When there exist one LacI molecule and one *lac*-promoter within an *E. coli* cell, the search time is less than 6 min = 0.1 hr [21]. In the case of *N_a_* unoccupied *lac*-promoters and *N_lacI_* copies of *LacI*, the search time becomes 0.1*/N_lacI_ Na*, which is equal to the inverse of the inactivation rate (1*/k_off_ N_a_*). Therefore, *k_off_* is estimated to be 10*N_lacI_* hr^−1^. *N_lacI_* can be roughly estimated from the fact that the strength of the lacI^*q*^ is similar to the promoter J23101 (*α_m_* of J23101 is 110 hr^−1^) [32]. Thus, the genomic expression level of LacI, *N_lacI_*, becomes *α_p_*[mRNA]_*lacI*_ /*γ_p_* =*α_p_α_m_/γ_p_γ_m_* = 1400 *·* 110*/*1 *·* 30 ≃ 5100. Thus, *k_off_* can be roughly estimated as 5.1 *×* 104 hr ^1^.

Gene activation, 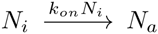: The activation rate constant *k_on_* is related to how fast the genomically-expressed LacI detached from its specific promoter *plac* (BBa_R0010). Considering that the dissociation constant is in the range of 0.1 − 1 pM = 10^−4^ 10^−3^ (copy number unit; here we used 1 nM corresponds to roughly 1 molecule number in the volume of *E. coli*) [34], [35], *k_on_* can be in the range of *k_off_ ×* (10^−4^ −10^−3^) = 5.1 − 51 hr^−1^.

#### E. Noise level correction

The mean level was corrected with a simple subtraction. The noise level was corrected by using the property that the observed variance (Variance*o*) is the sum of the GFP variance (Variance*g*) and the background signal variance (Variance*b*) under the assumption that the GFP signals are statistically independent of the background signals. More precisely, the noise level of GFP signals, defined by the square coefficient of variation, can be obtained by

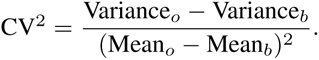

where the subscripts *o* and *b* denote observed and background signals, respectively.

#### F. Fluorescence histogram correction

The effect of autofluorescence was removed from the GFP signal histogram, more precisely probability mass function (pmf), by assuming that the autofluorescence is statistically independent of the true GFP signals [36]. Under this assumption, the pmf of the measured GFP signals, *T* (*v*), is related to both the autofluorescence pmf *v*(*v*) and the true GFP signal pmf *S*(*v*) via convolution:

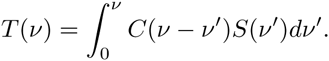

*S*(*v*) is obtained by minimizing the fitness function:

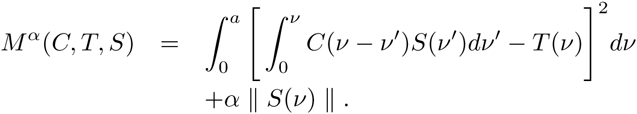

Here, *a* is the value of *v*beyond which *T* (*v*) is essentially zero, and in our study, we used the entire range of pmf. ‖ *S*(*v*) ‖ is a regularization term, defined as

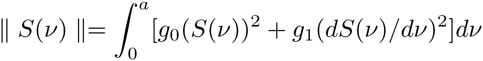

where *g*_0_ and *g*_1_ are positive regularization constants. The optimized solution of *S*(*v*) is obtained by following the procedure described below.

1. Remove background noise that is equipment-specific. Fluorescence signals of strength 0 and 1 (Sony Biotechnology ec800) were considered as background noise and removed. Then, *T* (*v*) (for the induction case of [IPTG] *>* 0) and *C*(*v*) (for the case of [IPTG]=0) were computed from the fluorescence signals using 1000 equal-width bins to obtain individual bin-sizes. Here, the bin-size of *T* is larger than that of *C*.
2. To compute the convolution, we will set the bin-size of *C* equal to that of *T*. Compute *C* again from the raw data using the bin-size of *T*, and append an array of zero at the end of *C* to make the total bin number equal to 1000.
3. Set the initial values of *S* equal to *T*.
4. Generate two different random numbers *v*_1_ and *v*_2_ in the range of [0, 999]. *S*(*v*_1_) and *S*(*v*_2_) were added and subtracted, respectively, by a constant *dS* = 10^−6^: *S*(*v*_1_) *S*(*v*_1_) + *dS* and *S*(*v*_2_) *S*(*v*_2_) *dS*. When *S*(*v*_2_) *dS* is less than zero, set *S*(*v*_1_) equal to *S*(*v*_1_) + *S*(*v*_2_) and then *S*(*v*_2_) equal to 0. In this way, the new *S* is automatically re-normalized and guaranteed to be non-negative.
5. Compute *M ^α^*. If *M ^α^* decreases, we accept the change and, otherwise, reject it and revert *S*(*v*) to the old *S* values before the update.
6. Repeat the steps 4 and 5 until *M ^α^* converges and compare 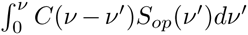 and *T* (*v*), where *S_op_* is the obtained optimized solution of *S*. If *S_op_*(*v*) shows oscillation, reduce the value of *α* while rebalancing *g*0 and *g*1 and go back to the step 4. If *S_op_* is noisy, increase the value of *α* while rebalancing *g*_0_ and *g*_1_ and go back to the step 4.

**TABLE I.** CONSTANTS USED IN THE AUTOFLUORESCENCE COMPENSATION ALGORITHM

| Constants | Values |
| --- | --- |
| $a$ | 1000 |
| $\alpha$ | 0.5 |
| $g_0$ | 0.001 |
| $g_1$ | 1.0 |
| $dS$ | $10^{-6}$ |

The constants used for the optimization are listed in Table I. The analysis was performed with Python 2.7.9 with Numpy 1.9.2, Scipy 0.15.1, and Spyder 2.3.4. Our python code is provided in the Supplementary Notes.

#### G. Nonlinear Regression

The gamma distribution function was used to fit our flow cytometry data. Protein copy number *N*_*pr*_ can be converted to fluorescence signal intensity *x*: *N*_*pr*_ = *c*_*s*_*x* with *c*_*s*_ a scaling constant. The gamma distribution function can be rescaled:

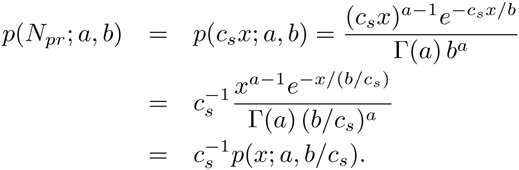

Here, Γ is a gamma function, and

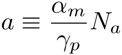

is the number of mRNA produced per cell doubling time, called burst frequency with *N_a_* the number of active promoters, and

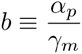

is the number of proteins produced during the mRNA lifetime, called burst size. Therefore, the fluorescence intensity should also follow the gamma distribution if its corresponding copy number follows the gamma distribution, with the burst size rescaled with *c_s_*. Nonlinear regression was carried by using the Scipy curve fit function (http://www.scipy.org/), which employs the Levenberg-Marquardt algorithm for the least squares fitting to estimate *a* and *b*.

## ACKNOWLEDGMENT

This work was supported by the National Science Foundations (NSF MCB 1158573).

## Supplementary Notes

### 1 DNA sequences of promoter-RBS-insertion-start codon regions

Table 1 shows the DNA sequences that were used for the region of ribosome binding sites along with different kinds of spacer sequences:

**Table 1:**
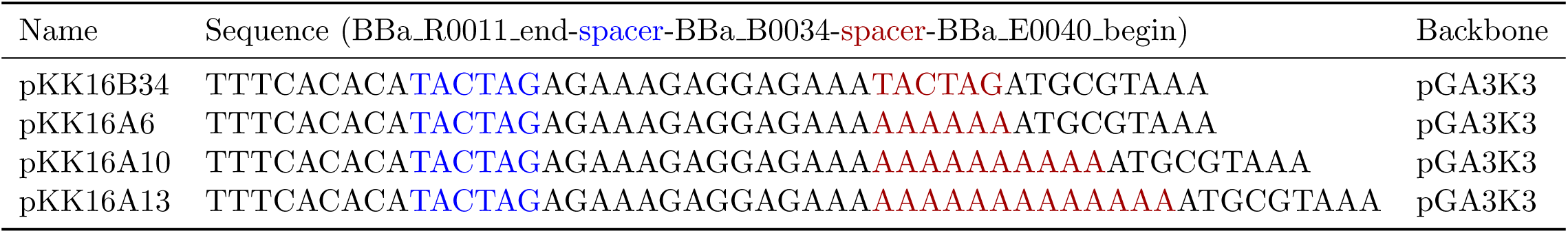
RBS region DNA sequences

### 2 Autofluorescence compensation in fluorescence histograms

Depending on the value of the regularization constant *α*, the optimized solution *S*(*v*) can show oscillation. In this section, we provide its sample pictures. There is a trade-off between the strength of noise in *S* and the fitting accuracy (comparison between the black and the blue dots in Fig. 2).

The python code used for the compensation is provided below.

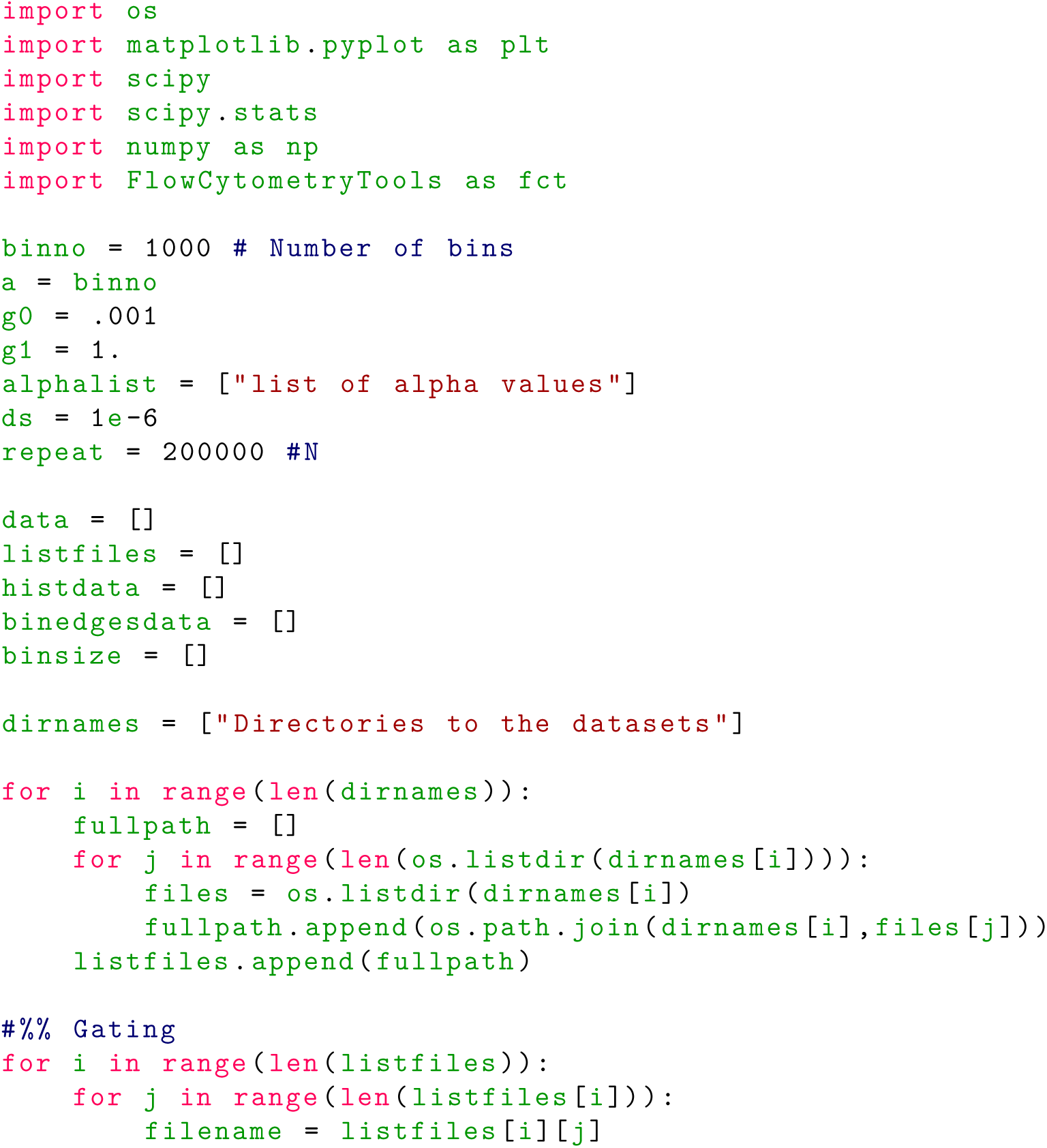
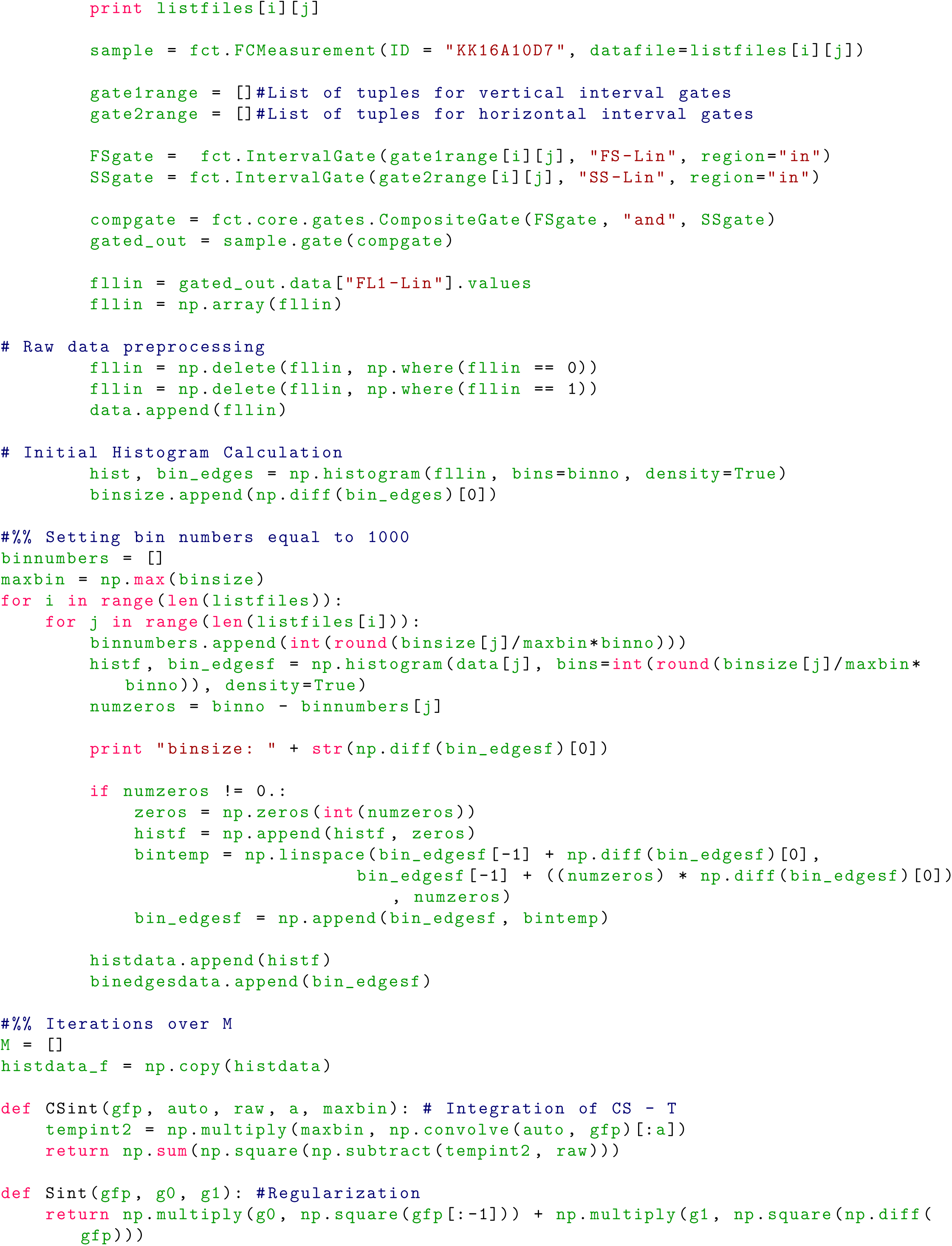
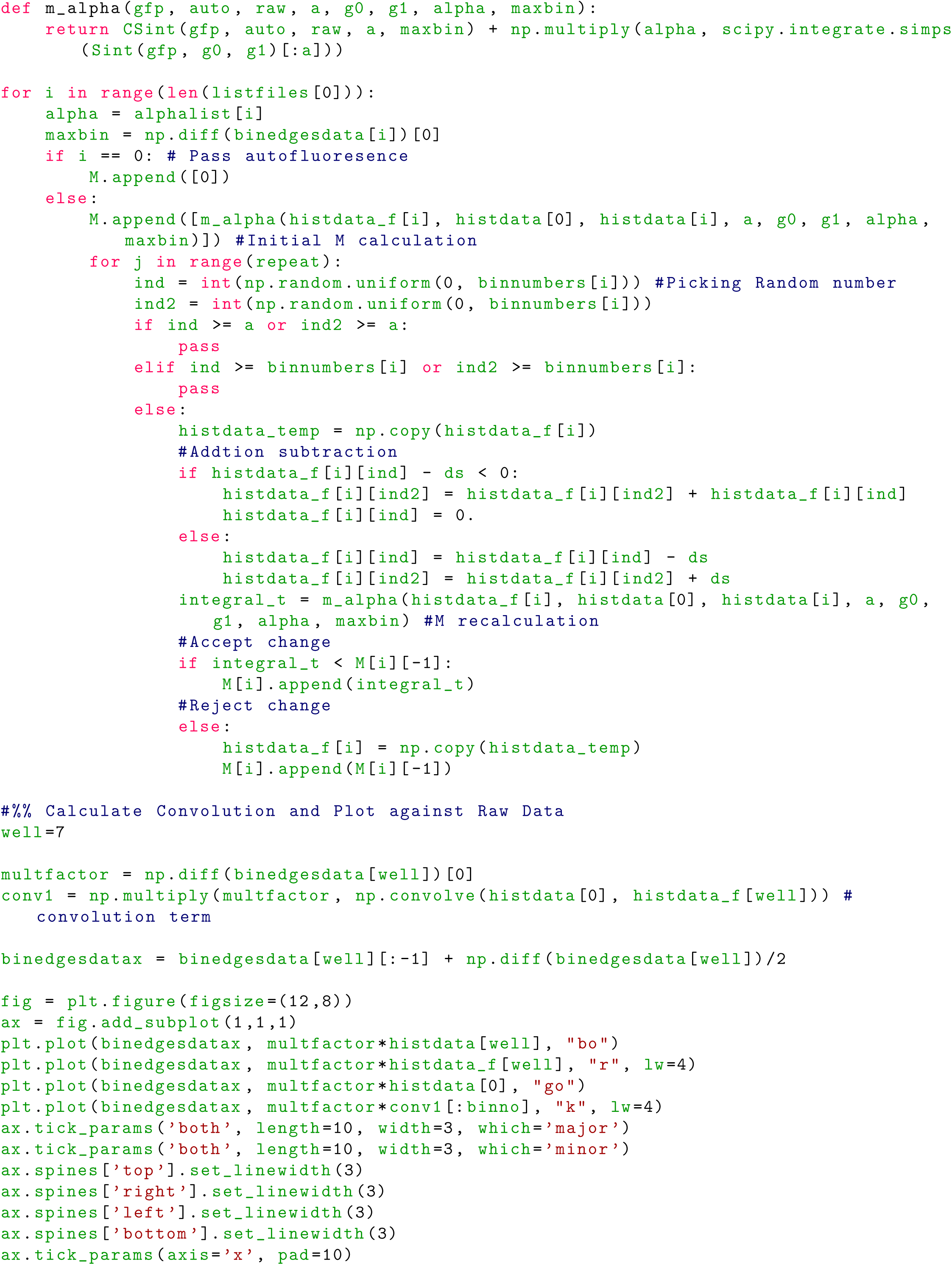
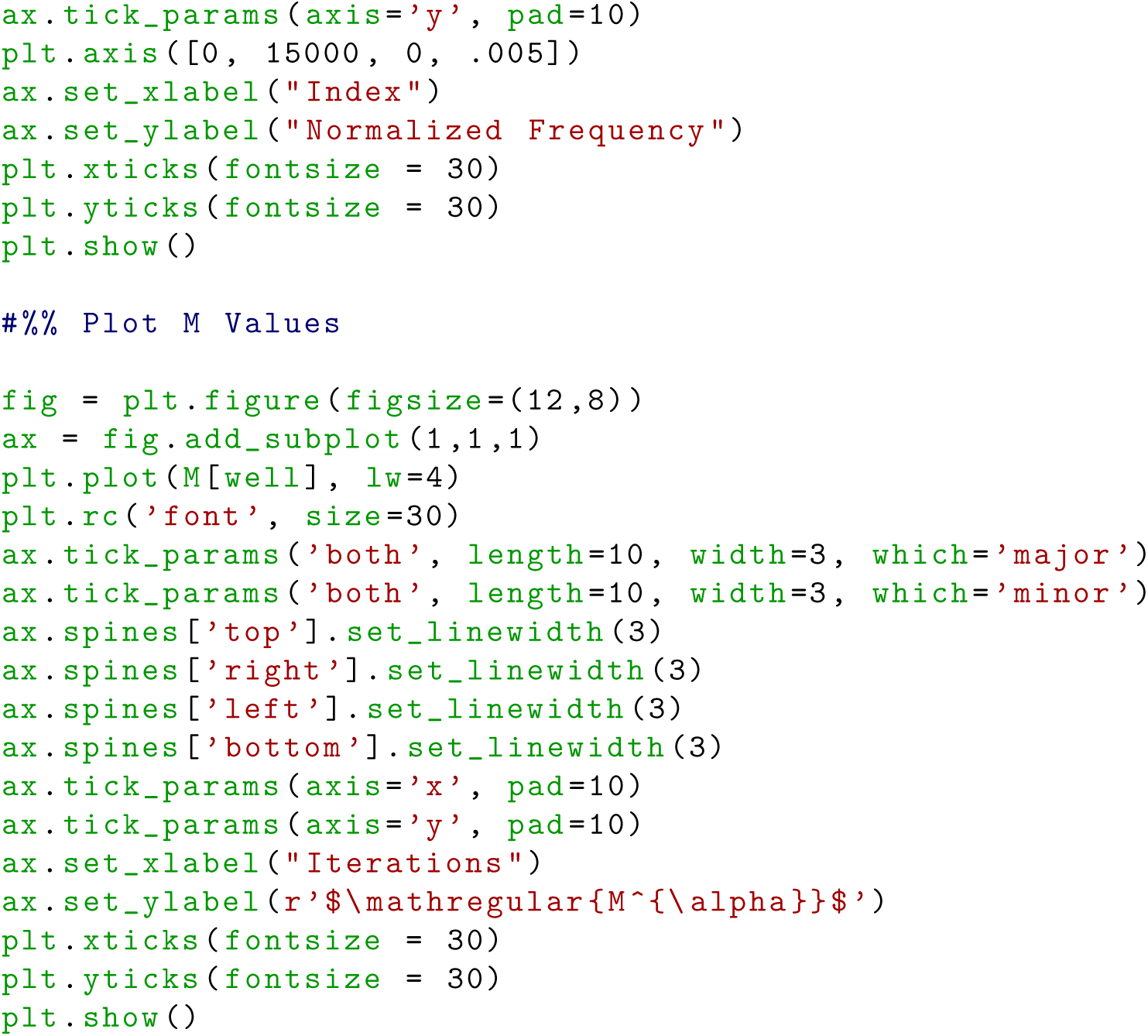

**Figure 1:**
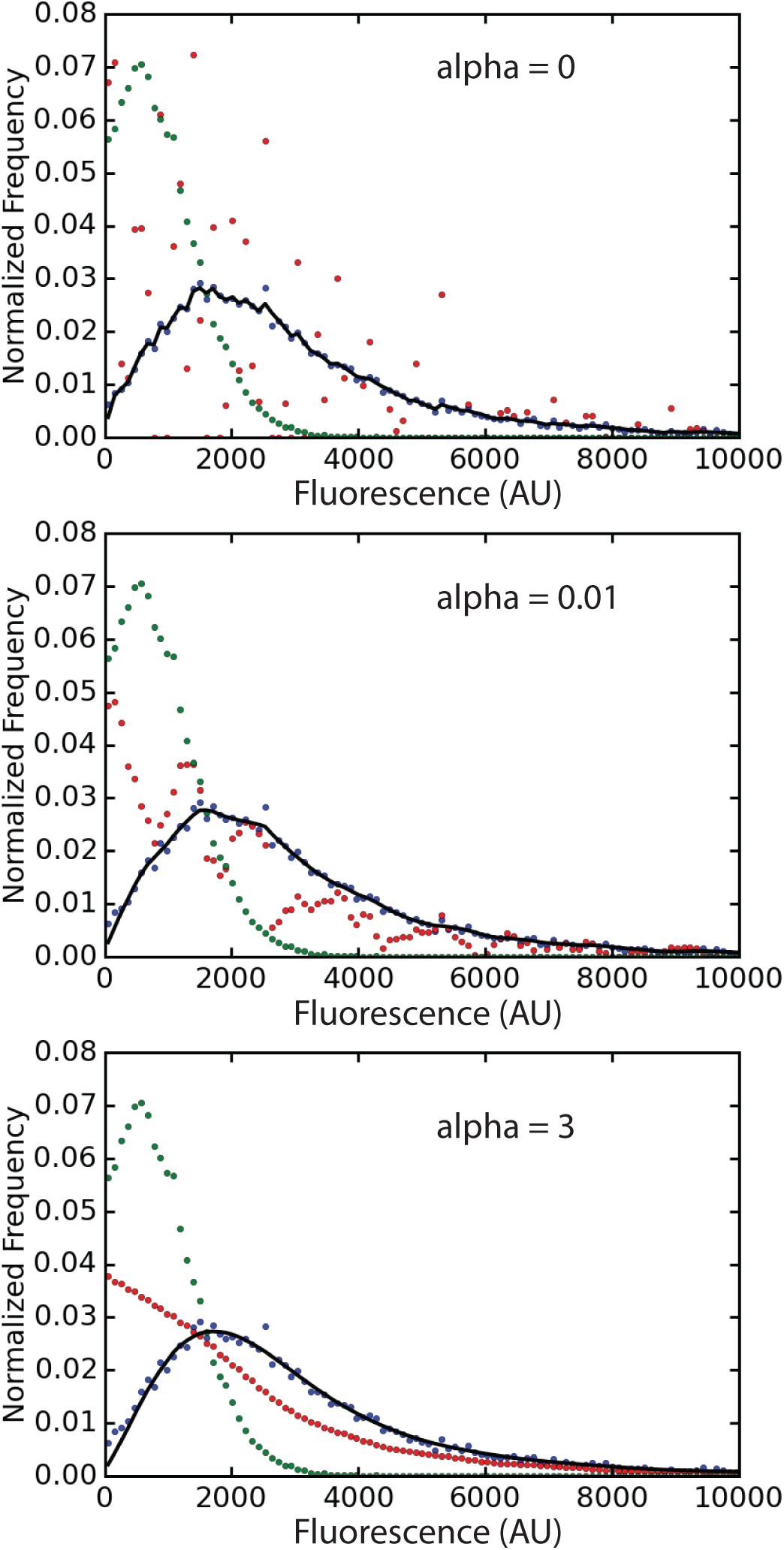
Autofluorescence compensation: The green dots correspond to autofluorescence (IPTG=0 case), the blue dots to the measured GFP signals, and the red dots to the optimized solution *S*, i.e., the true GFP signal probability mass function (normalized frequency). The black line is to verify the optimized solution *S* can generate *T* via convolution (refer to the Materials and Methods in the main manuscript).

**Figure 2:**
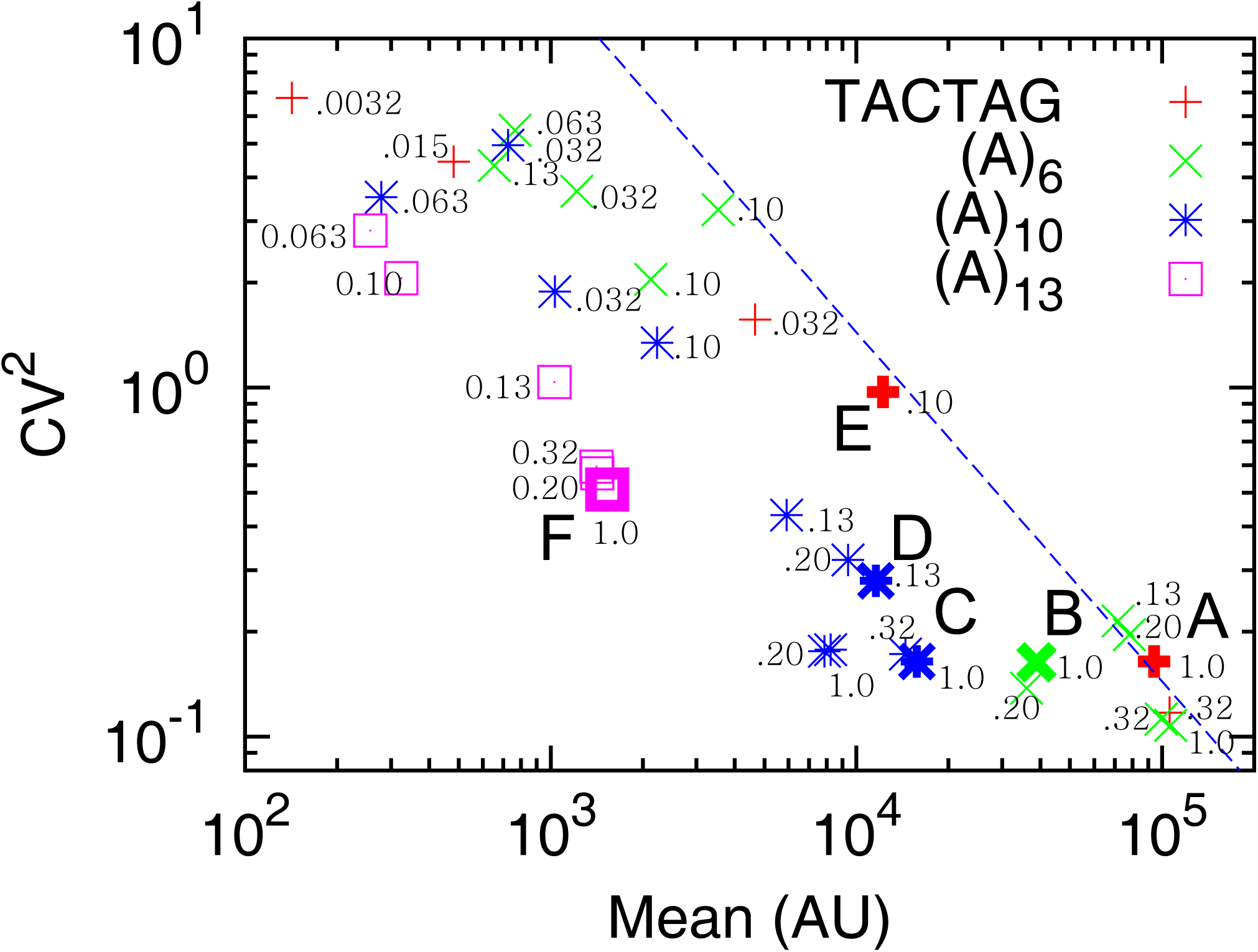
This figure is exactly the same as Figure 3 in the main manuscript. IPTG concentrations are included in the unit of mM. For (*A*)_6_ and (*A*)_10_, two biological replicates were used.

